## Supplementary Tables S1-S2 and Figures S1-S14 for "An Arg/Ala-Rich Helix in the N-Terminal Region of *M. tuberculosis* FtsQ Anchors FtsZ to Membranes"

*Supplementary Material*

Table S1. Chemical shifts assignments of FtsQ1-99

| Residue number | Residue name | N | HN | CO | Cα | Cβ |
|---|---|---|---|---|---|---|
| 1 | M | 119.1 | 8.346 | 176.4 | 55.54 | 32.52 |
| 2 | T | 114.5 | 8.011 | 174.4 | 61.75 | 69.64 |
| 3 | E | 122.8 | 8.319 | | | |
| 4 | H | 119.9 | 8.352 | | | |
| 5 | N | 121.0 | 8.390 | 174.7 | 52.99 | 39.06 |
| 6 | E | 121.3 | 8.456 | 175.7 | 56.09 | 30.19 |
| 7 | D | 122.9 | 8.370 | | | 40.76 |
| 8 | P | | | 177.0 | 63.21 | 31.90 |
| 9 | Q | 119.8 | 8.492 | 175.9 | 55.72 | 28.86 |
| 10 | I | 121.2 | 7.916 | 175.9 | 61.02 | 38.57 |
| 11 | E | 124.8 | 8.367 | 176.0 | 56.37 | 30.10 |
| 12 | R | 123.0 | 8.285 | 176.0 | 55.68 | 30.78 |
| 13 | V | 122.1 | 8.209 | 175.7 | 61.99 | 32.54 |
| 14 | A | 127.8 | 8.368 | 177.3 | 52.30 | 19.21 |
| 15 | D | 120.1 | 8.266 | 176.0 | 54.24 | 41.04 |
| 16 | D | 120.5 | 8.223 | 176.0 | 54.12 | 40.93 |
| 17 | A | 123.9 | 8.105 | 177.5 | 52.48 | 18.97 |
| 18 | A | 122.8 | 8.136 | 177.5 | 52.47 | 19.02 |
| 19 | D | 119.5 | 8.188 | 176.4 | 54.29 | 41.03 |
| 20 | E | 121.2 | 8.277 | 176.5 | 56.71 | 30.20 |
| 21 | E | 121.2 | 8.322 | 176.1 | 56.42 | 30.04 |
| 22 | A | 124.8 | 8.151 | 177.5 | 52.32 | 18.98 |
| 23 | V | 119.2 | 8.056 | 176.2 | 62.11 | 32.51 |
| 24 | T | 117.8 | 8.140 | 174.1 | 61.49 | 69.74 |
| 25 | E | 124.6 | 8.279 | 174.2 | 54.20 | 29.68 |
| 26 | P | | | 176.7 | 62.97 | 31.80 |
| 27 | L | 122.0 | 8.303 | 177.3 | 54.93 | 42.22 |
| 28 | A | 125.1 | 8.306 | 177.8 | 52.45 | 18.92 |
| 29 | T | 112.6 | 7.991 | 174.5 | 61.74 | 69.56 |
| 30 | E | 123.0 | 8.332 | 176.2 | 56.43 | 30.16 |
| 31 | S | 117.1 | 8.325 | 174.4 | 58.08 | 63.60 |
| 32 | K | 123.4 | 8.371 | 176.0 | 56.15 | 32.78 |
| 33 | D | 120.9 | 8.258 | 175.6 | 54.04 | 41.16 |
| 34 | E | 122.1 | 8.136 | 174.3 | 54.11 | 29.75 |
| 35 | P | | | 176.6 | 63.08 | 31.80 |
| 36 | A | 123.9 | 8.330 | 177.5 | 52.27 | 19.16 |
| 37 | E | 119.3 | 8.233 | 175.8 | 56.38 | 30.26 |
| 38 | H | 121.7 | 8.349 | 176.1 | 56.41 | 30.09 |
| 39 | P | | | 177.1 | 63.67 | 31.86 |
| 40 | E | 120.4 | 9.091 | 176.4 | 56.98 | 29.43 |
| 41 | F | 119.8 | 8.006 | 175.4 | 57.12 | 39.40 |
| 42 | E | 122.2 | 8.184 | 176.3 | 56.49 | 30.53 |
| 43 | G | 110.1 | 8.156 | 172.4 | 45.20 | |
| 44 | P | | | 178.2 | | |
| 45 | R | 120.3 | 8.315 | | | |
| 46 | R | 119.8 | 8.292 | | | |
| 47 | R | 121.4 | 8.249 | | | |
| 48 | A | 123.3 | 8.153 | 179.0 | 53.61 | |
| 49 | R | 119.5 | 8.091 | 177.6 | | |
| 50 | R | 121.9 | 8.232 | 177.3 | 57.58 | 30.08 |
| 51 | E | 120.8 | 8.466 | 178.0 | 58.02 | 29.71 |
| 52 | R | 120.5 | 8.193 | 177.5 | 57.94 | 30.23 |
| 53 | A | 123.7 | 8.126 | 179.6 | 53.96 | 18.38 |
| 54 | E | 119.9 | 8.445 | 178.3 | 57.93 | 29.50 |
| 55 | R | 121.4 | 8.187 | 178.0 | 57.93 | 29.66 |
| 56 | R | 120.7 | 8.195 | 178.0 | 57.81 | 29.95 |
| 57 | A | 123.6 | 8.172 | 179.2 | 53.78 | 18.28 |
| 58 | A | 122.0 | 8.138 | 179.7 | 53.93 | 18.34 |
| 59 | Q | 119.2 | 8.165 | 177.3 | 57.21 | 28.60 |
| 60 | A | 123.8 | 8.144 | 179.4 | 53.86 | 18.28 |
| 61 | R | 119.6 | 8.080 | 177.3 | 57.60 | 30.22 |
| 62 | A | 122.9 | 8.067 | 179.7 | 53.96 | 18.37 |

| | | | | | | |
|---|---|---|---|---|---|---|
| 63 | T | 114.4 | 8.241 | 175.4 | 64.23 | 69.23 |
| 64 | A | 125.0 | 8.088 | 180.3 | 54.47 | 18.15 |
| 65 | I | 119.9 | 8.037 | 177.9 | 63.62 | 38.00 |
| 66 | E | 122.1 | 7.984 | 178.7 | 58.47 | 29.13 |
| 67 | Q | 119.2 | 8.413 | 178.2 | 58.14 | 27.55 |
| 68 | A | 122.9 | 8.005 | 179.9 | 54.20 | 18.02 |
| 69 | R | 119.8 | 8.122 | 178.4 | 58.28 | 30.10 |
| 70 | R | 119.9 | 8.080 | 177.9 | 58.14 | 30.16 |
| 71 | A | 122.7 | 8.040 | 178.8 | 53.69 | 18.35 |
| 72 | A | 121.4 | 7.921 | 179.0 | 53.55 | 18.42 |
| 73 | K | 119.2 | 7.905 | 177.4 | 57.21 | 32.46 |
| 74 | R | 120.4 | 7.966 | 177.0 | 57.03 | 30.42 |
| 75 | R | 121.1 | 8.109 | 176.5 | 56.68 | 30.55 |
| 76 | A | 124.3 | 8.092 | 177.9 | 52.56 | 18.89 |
| 77 | R | 119.6 | 8.154 | 176.9 | 56.44 | 30.46 |
| 78 | G | 109.1 | 8.284 | 173.9 | 45.13 | |
| 79 | Q | 119.8 | 8.104 | 175.7 | 55.58 | 29.41 |
| 80 | I | 122.5 | 8.196 | 176.3 | 61.01 | 38.35 |
| 81 | V | 124.8 | 8.250 | 175.9 | 62.10 | 32.57 |
| 82 | S | 119.6 | 8.317 | 174.4 | 58.09 | 63.74 |
| 83 | E | 123.1 | 8.425 | 176.2 | 56.58 | 30.12 |
| 84 | Q | 121.0 | 8.323 | 175.3 | 55.67 | 29.29 |
| 85 | N | 120.5 | 8.446 | 173.0 | | |
| 86 | P | | | 176.5 | 63.11 | 31.87 |
| 87 | A | 123.4 | 8.231 | 177.4 | 52.13 | 18.91 |
| 88 | K | 121.4 | 8.086 | 174.2 | 53.86 | 32.38 |
| 89 | P | | | 176.6 | 62.95 | 31.86 |
| 90 | A | 124.4 | 8.348 | 177.4 | 52.12 | 19.02 |
| 91 | A | 123.6 | 8.231 | 177.3 | 52.21 | 19.13 |
| 92 | R | 120.3 | 8.270 | 176.7 | 56.05 | 30.64 |
| 93 | G | 110.0 | 8.332 | 173.7 | 45.07 | |
| 94 | V | 119.7 | 7.948 | 176.1 | 62.17 | 32.54 |
| 95 | V | 125.1 | 8.240 | 175.9 | 62.16 | 32.48 |
| 96 | R | 125.8 | 8.433 | 176.4 | 55.96 | 30.75 |
| 97 | G | 110.2 | 8.369 | 173.7 | 45.04 | |
| 98 | L | 122.1 | 8.078 | 176.4 | 55.19 | 42.37 |
| 99 | K | 126.4 | 7.808 | 181.0 | 57.47 | 33.62 |

Table S2. Differences in average chemical shifts between helical and nonhelical positions

| | Helical | | | Nonhelical[2] | | |
|---|---|---|---|---|---|---|
| **Residue type** | **$C^O$** | **$C_\alpha$** | **$C_\beta$** | **$C^O$** | **$C_\alpha$** | **$C_\beta$** |
| Ala (7[a] vs. 9[b]) | 179.7 ± 0.3[c] | 54.0 ± 0.2 | 18.3 ± 0.1 | 177.5 ± 0.1 | 52.3 ± 0.1 | 19.0 ± 0.1 |
| Arg (7[a] vs. 5[b]) | 177.8 ± 0.4 | 57.9 ± 0.2 | 30.1 ± 0.2 | 176.5 ± 0.3 | 56.2 ± 0.4 | 30.6 ± 0.1 |
| Glu (3[a] vs. 11[b]) | 178.3 ± 0.3 | 58.1 ± 0.2 | 29.4 ± 0.2 | 175.8 ± 0.8 | 56.1 ± 0.9 | 30.0 ± 0.3 |

[a]Number of this amino acid within the helical core, or residues 50-70.

[b]Number of this amino acid within nonhelical regions, or residues 1-42 and 80-99.

[c]Errors are standard deviations.

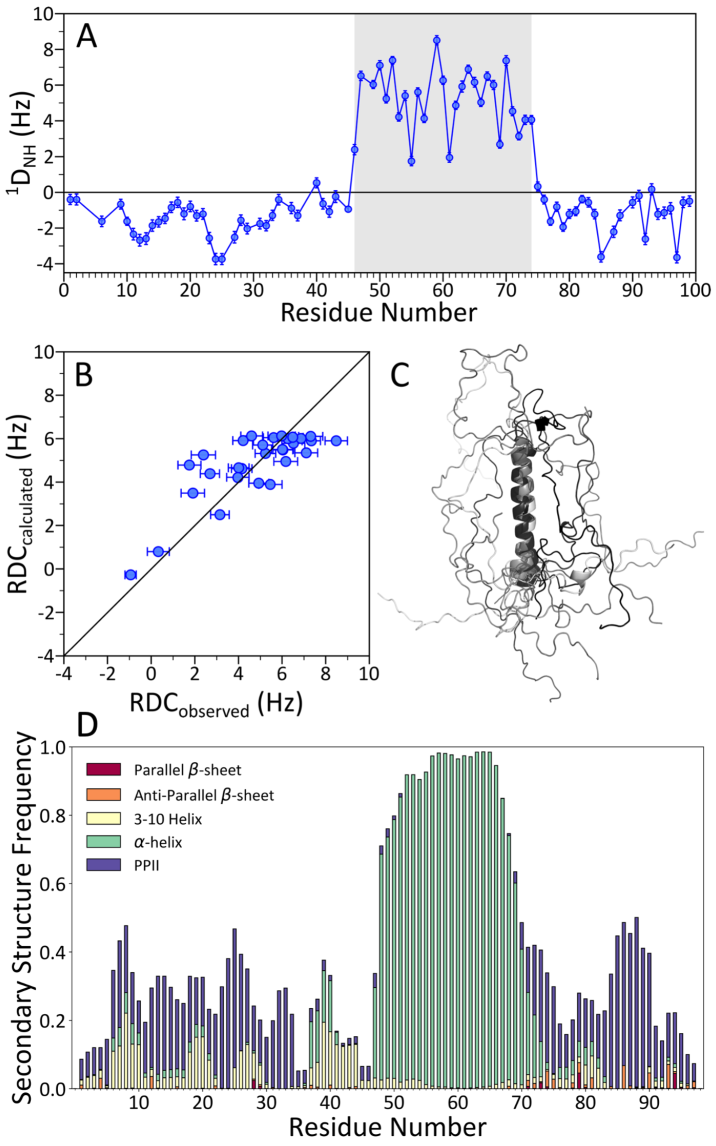


**Figure S1.** Helix formation in FtsQ1-99. (A) RDCs of uniformly $^{15}$N-labeled FtsQ1-99 aligned with neutral stretched acrylamide gels in 20 mM Tris (pH 6.85) with 50 mM NaCl measured at 800 MHz and 305 K. Helical residues (46-74) are highlighted in gray. (B) Correlation of observed RDC values for residues 45-75 with those back calculated from the lowest-energy model. (C) Eight lowest-energy models. (D) Secondary structure propensities in MD simulations.

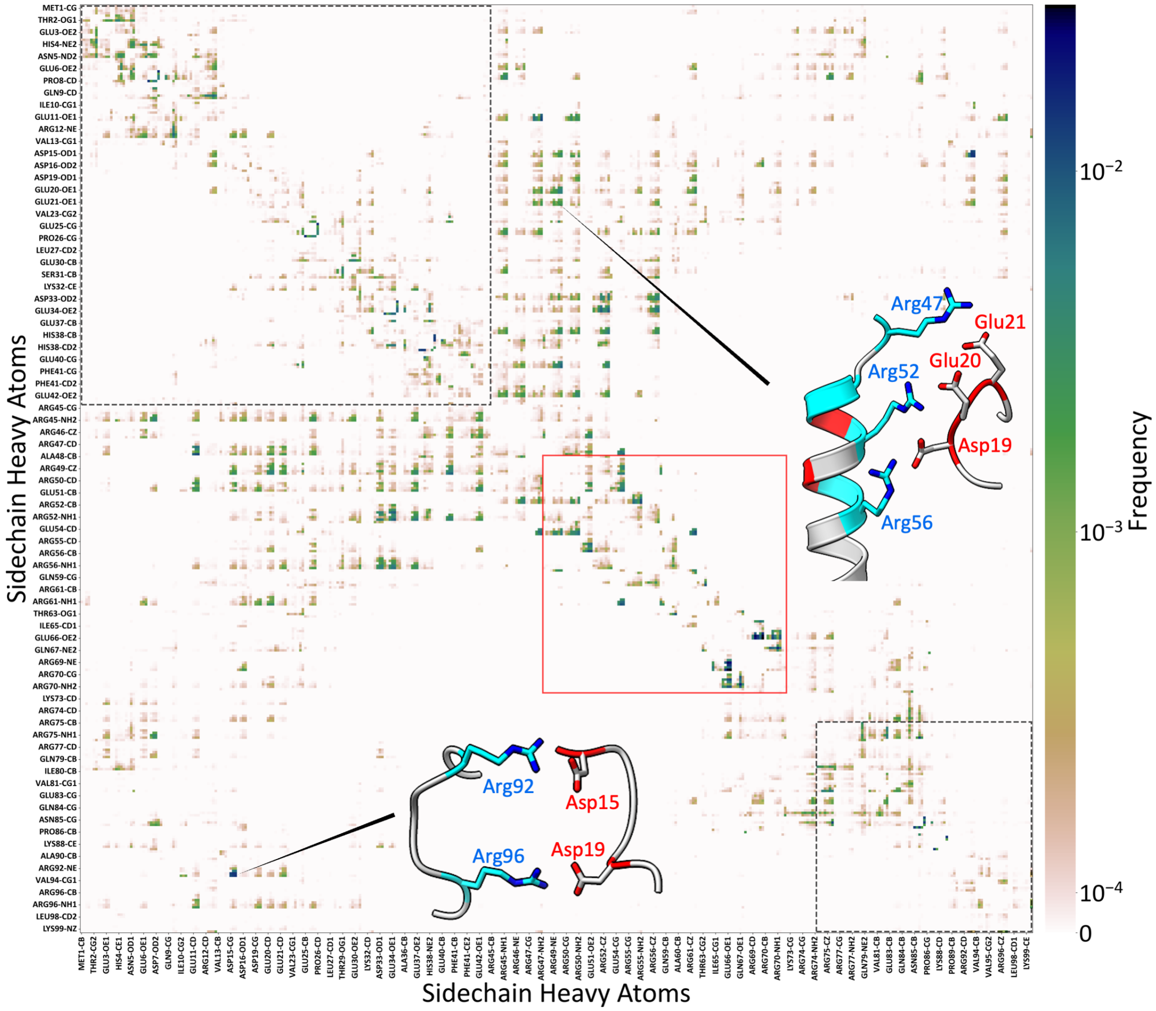


**Figure S2.** Sidechain-sidechain contact map of FtsQ1-99 from MD simulations in solution. The Arg/Ala-rich helix core (residues 48-72) is indicated by a red box; the N-tail (residues 1-44) and C-tail (residues 75-99) are indicated by a black dotted box. Inset: two snapshots illustrating the contact formation of the N-tail with either the Arg/Ala-rich helix or the C-tail.

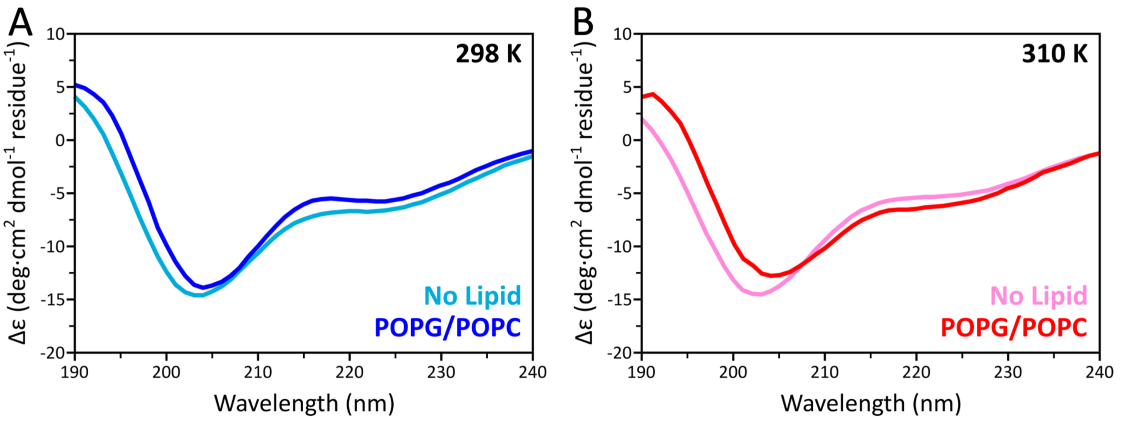


**Figure S3.** Circular dichroism spectra of 20 μM FtsQ1-99 in 2 mM Tris buffer (pH 6.85) with 5 mM NaCl measured at (A) 298 K and (B) 310 K, in the absence (marine and pink) and presence (blue and red) of POPG/POPC lipid vesicles (7:3 molar ratio). The protein-to-lipid was 1:16.5.

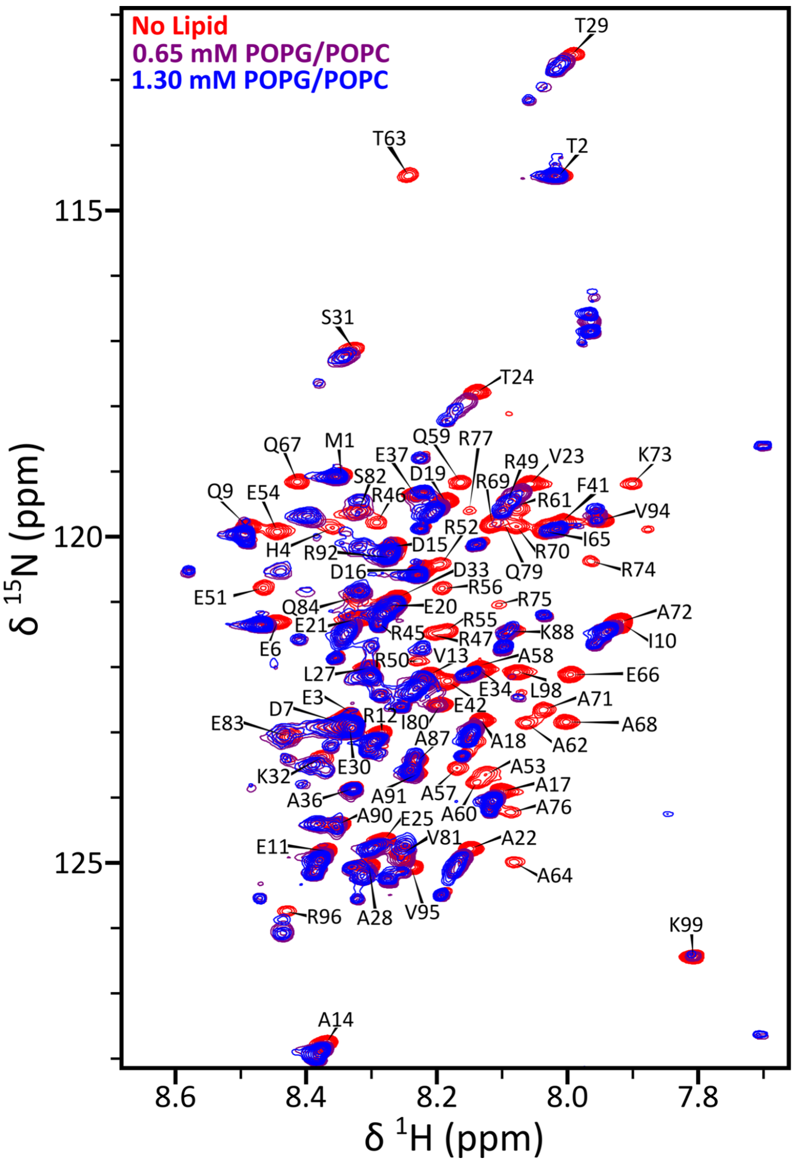


**Figure S4.** $^{1}H$-$^{15}N$ HSQC spectra of 130 μM uniformly $^{13}C$-$^{15}N$-labeled FtsQ1-99 in the absence (red) or presence of POPG/POPE (7:3) vesicles at 1:5 (purple) or 1:10 (blue) protein-to-lipid ratios. Acquired at 800 MHz and 305 K in 20 mM Tris buffer (pH 6.85) with 50 mM NaCl.

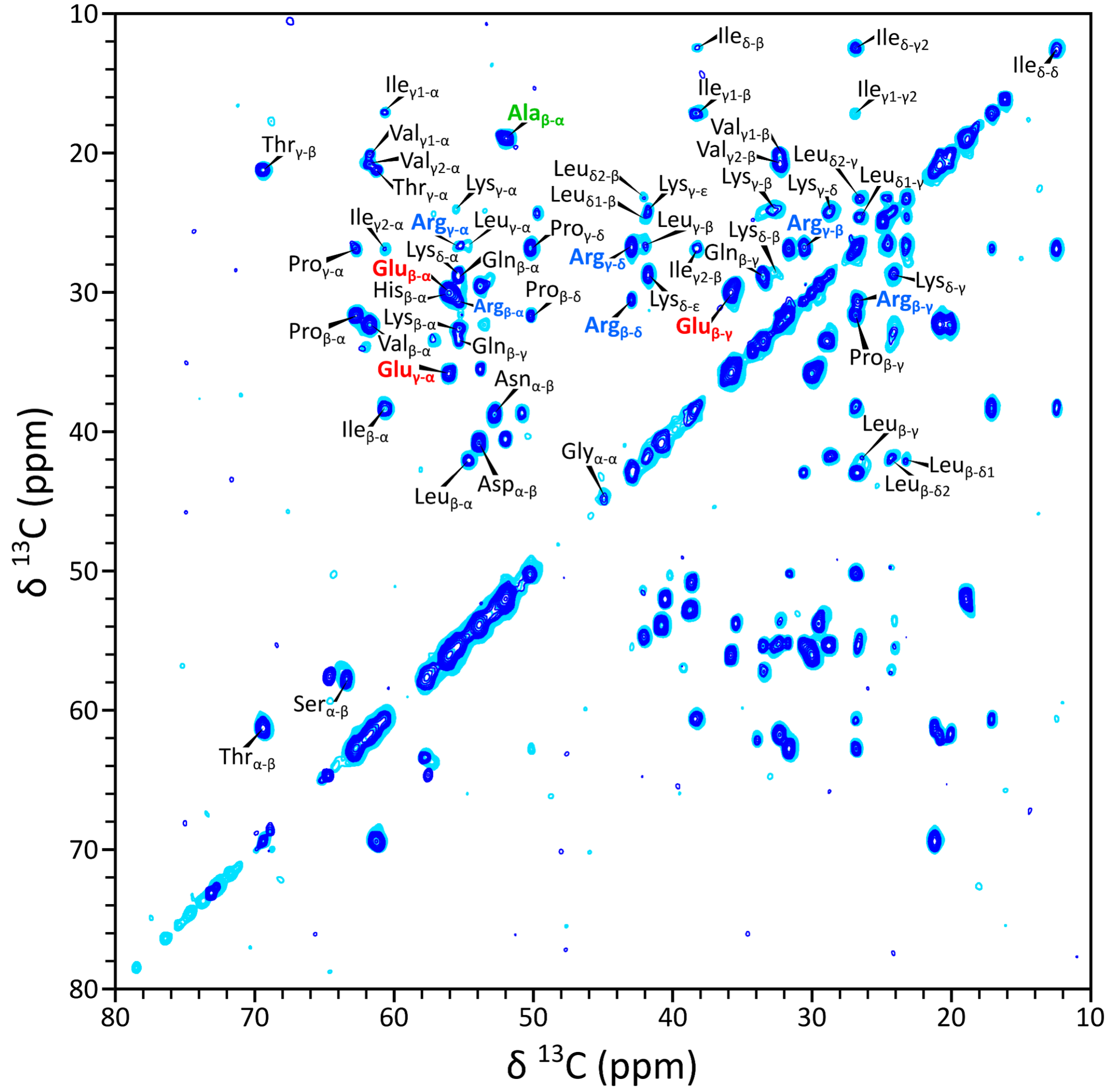


**Figure S5.** TOBSY-INEPT $^{13}C$-$^{13}C$ correlation spectra of FtsQ1-99 bound to 7:3 POPG/POPC (blue) or native (cyan) lipid vesicles. Acquired at 800 MHz with a 13 kHz spinning rate at 298 K in 20 mM Tris buffer (pH 6.85) with 50 mM NaCl.

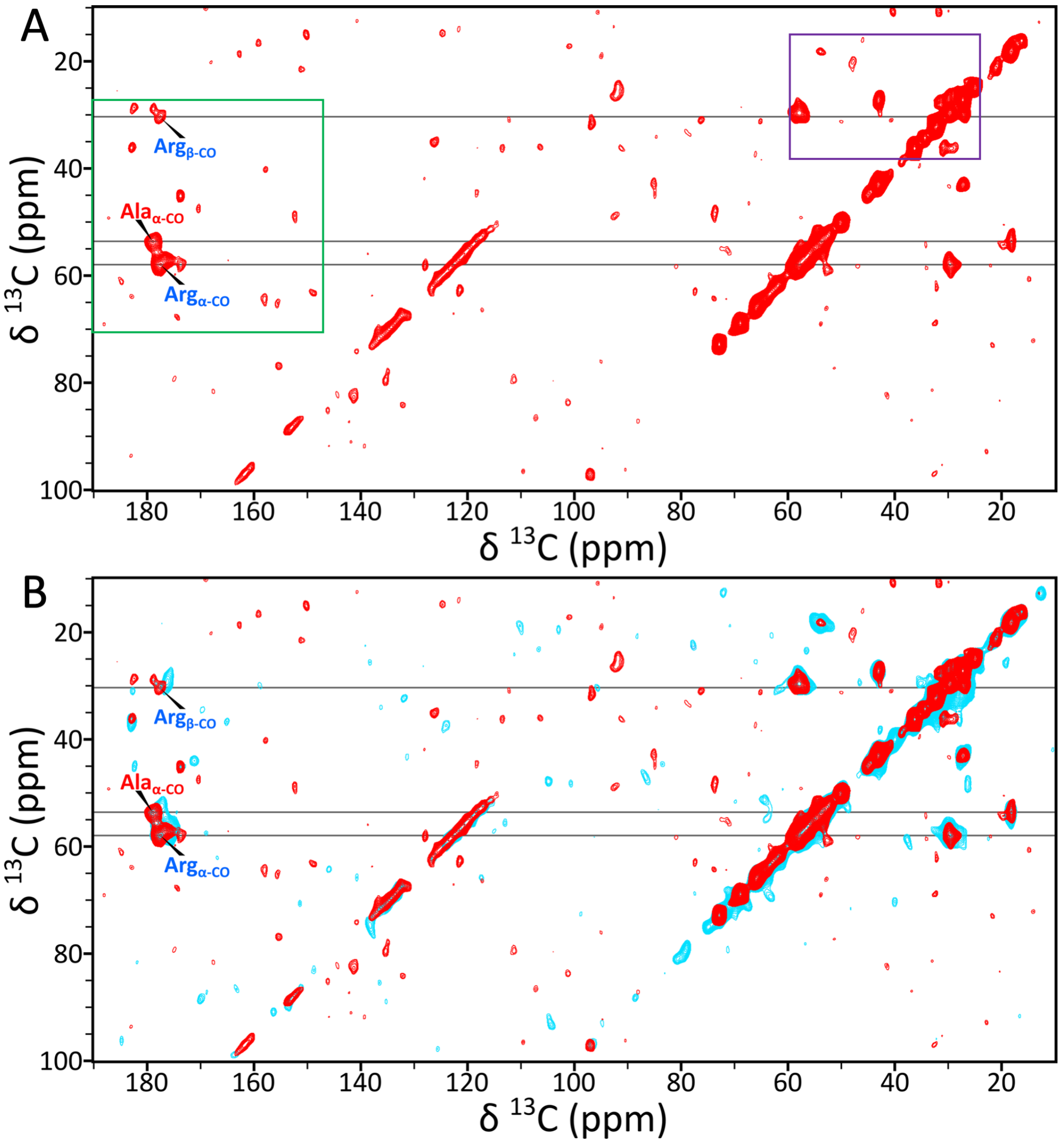


**Figure S6.** CP-PARIS $^{13}$C-$^{13}$C correlation spectra of FtsQ1-99 bound to lipid vesicles. (A) The spectrum of FtsQ1-99 bound to 7:3 POPG/POPC vesicles. The regions enclosed in the purple and green boxes are shown in Fig. 5A, B. (B) Overlay of this spectrum (red) with the counterpart using native lipid vesicles (cyan). Acquired at 800 MHz with a 13 kHz spinning rate at 285 K in 20 mM Tris buffer (pH 6.85) with 50 mM NaCl.

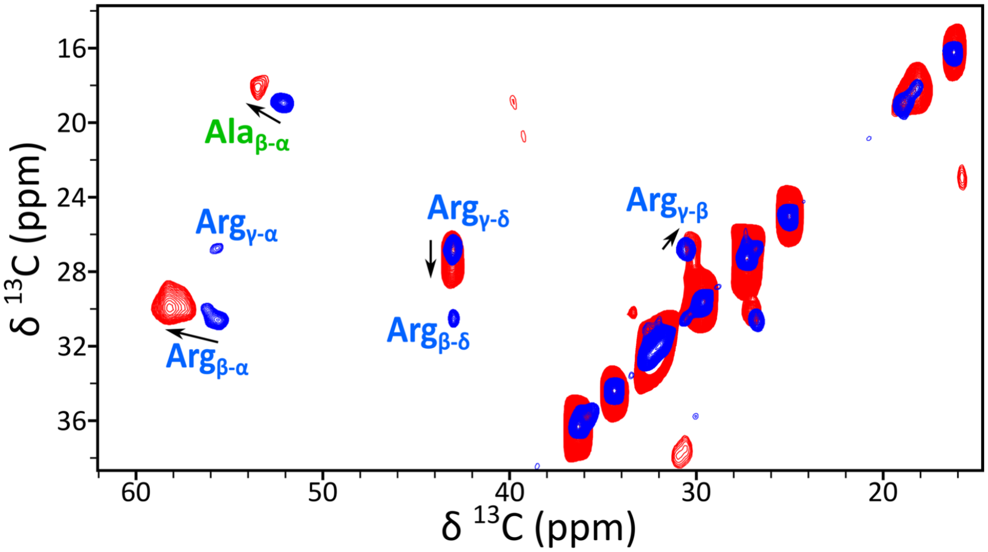


**Figure S7.** Identification of comparatively rigid and dynamic residues of FtsQ1-99 by MAS solid-state NMR using Ala and Arg specifically-labeled FtsQ1-99. Overlay of INEPT (blue) and CP (red) $^{13}$C-$^{13}$C correlation spectra. Acquired at 800 MHz with a 13 kHz spinning rate, employing 7.5 ms and 30 ms mixing times for TOBSY-INEPT (298 K) and CP-PARIS (285 K), respectively. The samples contained $^{13}$C$^{15}$N-[AR]-labeled FtsQ1-99 bound to 7:3 POPG/POPC vesicles at a protein-to-lipid ratio of 1:20 in 20 mM Tris buffer (pH 6.85) with 50 mM NaCl. The selected region highlights the only significant sidechain resonances, assigned to Ala and Arg residues.

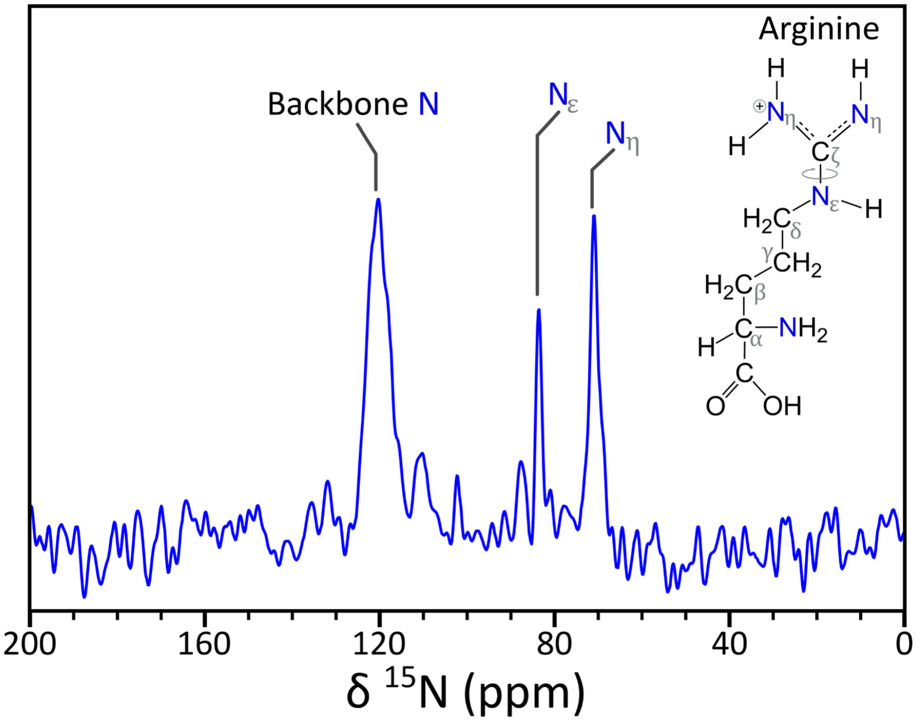


**Figure S8.** One-dimensional $^{15}N$ CP-MAS spectrum of uniformly $^{13}C^{15}N$-labeled FtsQ1-99 bound to POPG/POPC (7:3) vesicles in 20 mM Tris buffer (pH 6.85) with 50 mM NaCl. Acquired at 600 MHz and 295 K with a 8 kHz spinning rate.

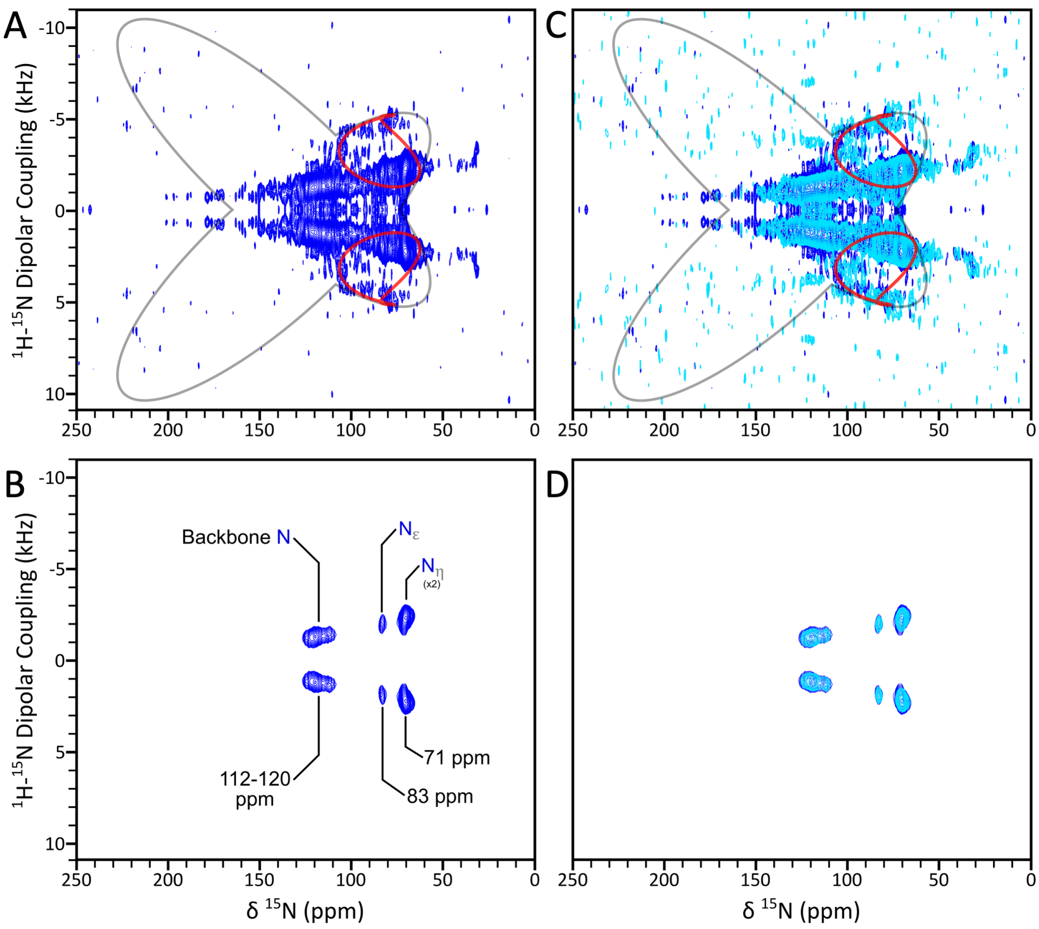


**Figure S9.** "Magic-sandwich" PISEMA spectra of uniformly $^{15}$N-labeled FtsQ1-99 bound to lipid bilayers mechanically oriented using glass slides. (A) Low-contour plot of the spectrum of FtsQ1-99 bound to 7:3 POPG/POPC bilayers, presenting a helical wheel with a lateral orientation relative to the bilayer surface. Also shown is a simulated powder pattern using average $^{15}$N CSA tensor parameters: $\sigma_{11}$ = 57.3 ppm, $\sigma_{22}$ = 81.2 ppm, and $\sigma_{33}$ = 228.1 (black) and a PISA helical wheel with a tilt of 74° (red). (B) High-contour plot highlighting the significant resonances assigned to Arg sidechain sites embedded in the membrane surface. The intensities spanning 112-120 ppm with modest dipolar alignment are likely from backbone nitrogens of residues flanking the helix. (C-D) Overlay with the counterparts using native lipid bilayers (cyan). Acquired at 800 MHz and 298 K on samples at a protein-to-lipid ratio of 1:40 in 5 mM Tris buffer (pH 6.85) with 2 mM NaCl.

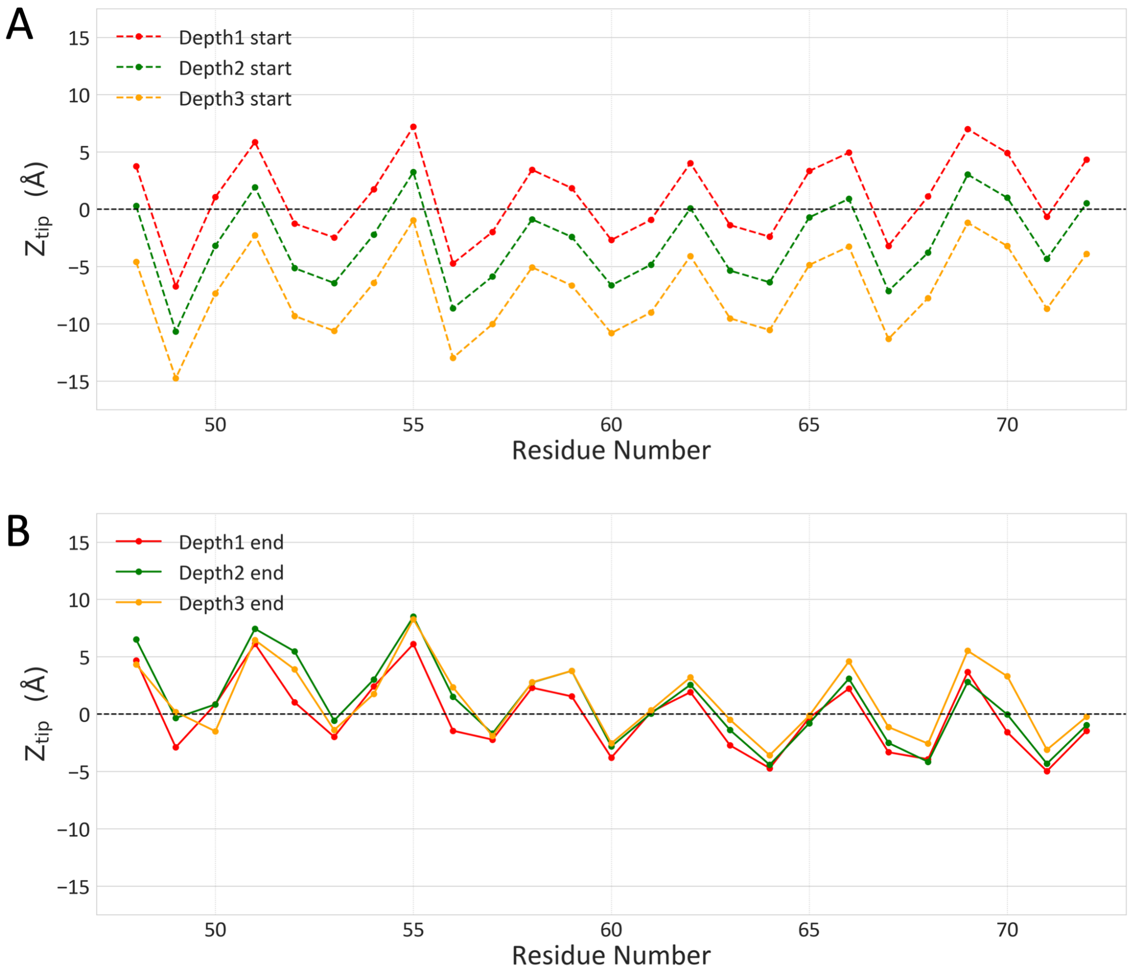


**Figure S10.** Burial depth of the Arg/Ala-rich helix in membranes. (A) $Z_{tip}$ distances of residues 48-72 sidechains in three initial structures for MD simulations. (B) Average $Z_{tip}$ distances in the last 10 ns of 200 ns MD simulations. The three simulations all converged to the same shallow burial depth.

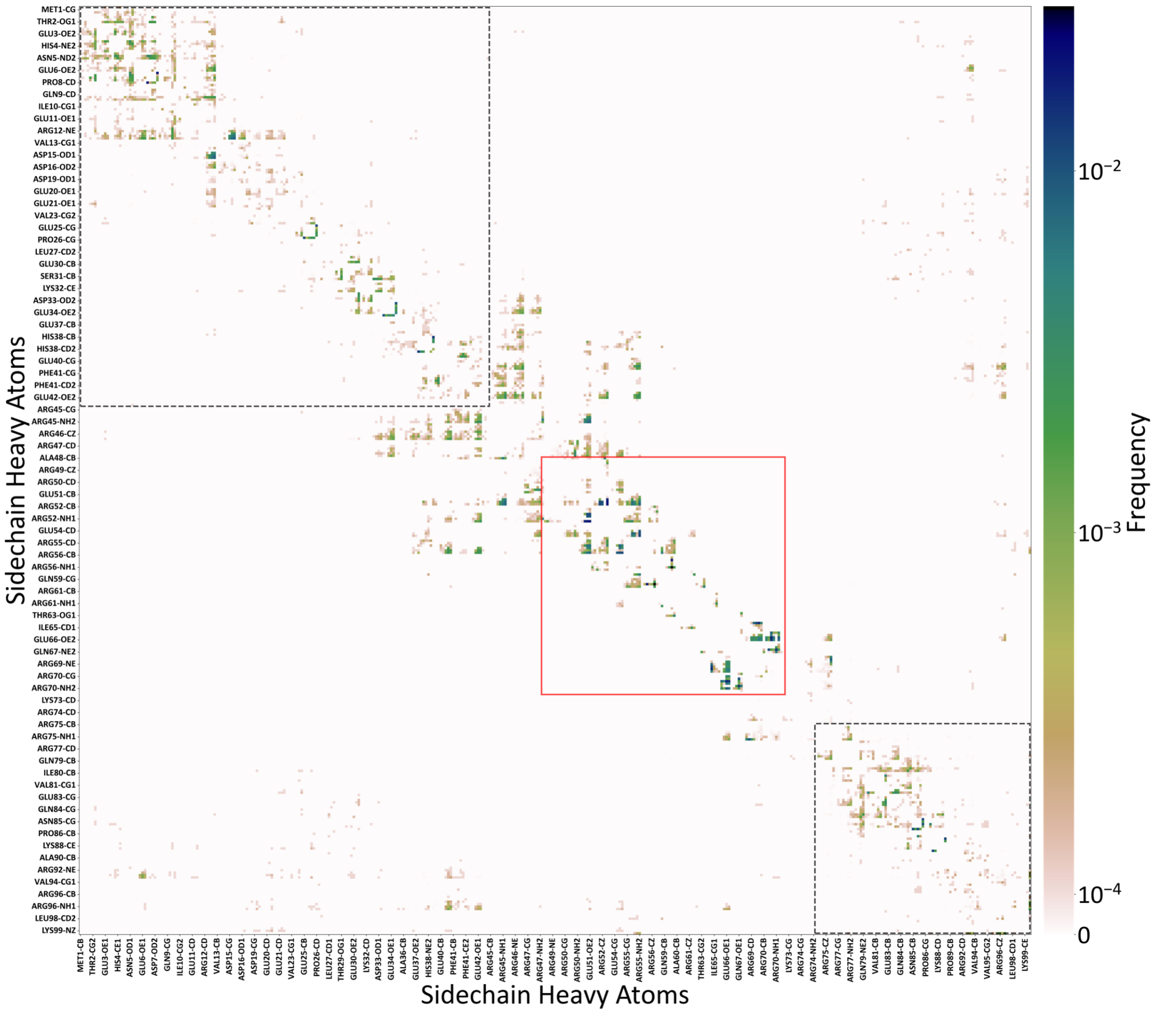


**Figure S11.** Sidechain-sidechain contact map of membrane-bound FtsQ1-99 from MD simulations. The Arg/Ala-rich helix core (residues 48-72) is indicated by a red box; the N-tail (residues 1-44) and C-tail (residues 75-99) are indicated by a black dotted box.

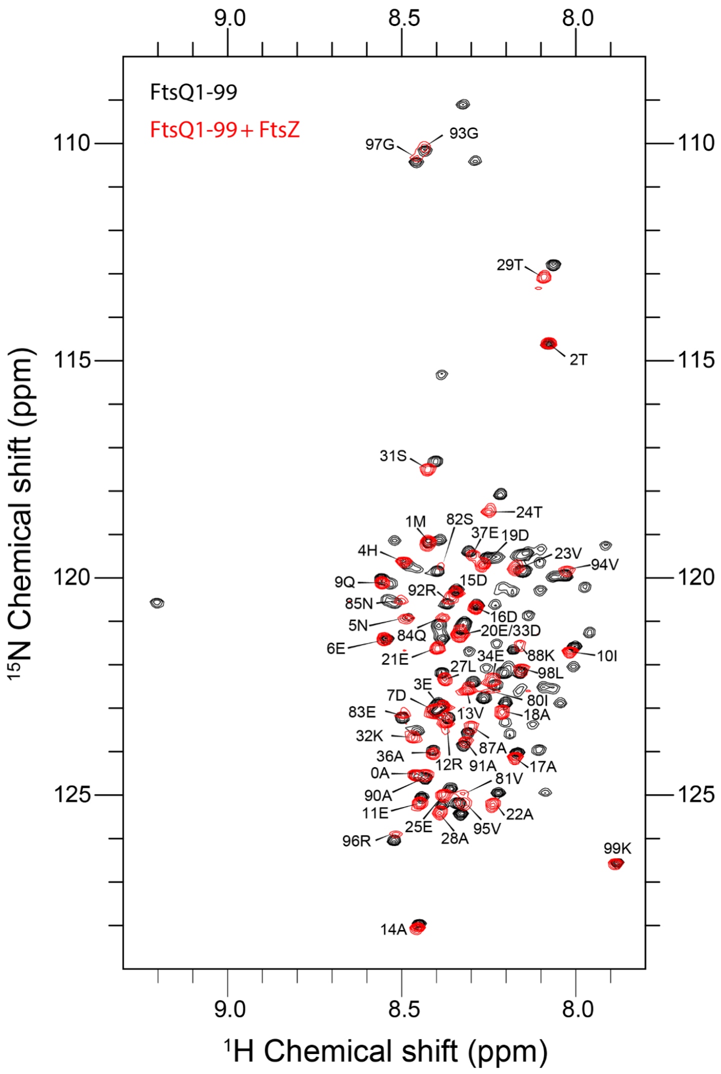


**Figure S12.** $^{1}H$-$^{15}N$ HSQC spectra of 50 μM uniformly $^{13}C$-$^{15}N$ labeled FtsQ1-99 in the absence (black) or presence (red) of 50 μM FtsZ. Acquired at 800 MHz 298 K in 20 mM phosphate buffer (pH 6.5) with 25 mM NaCl.

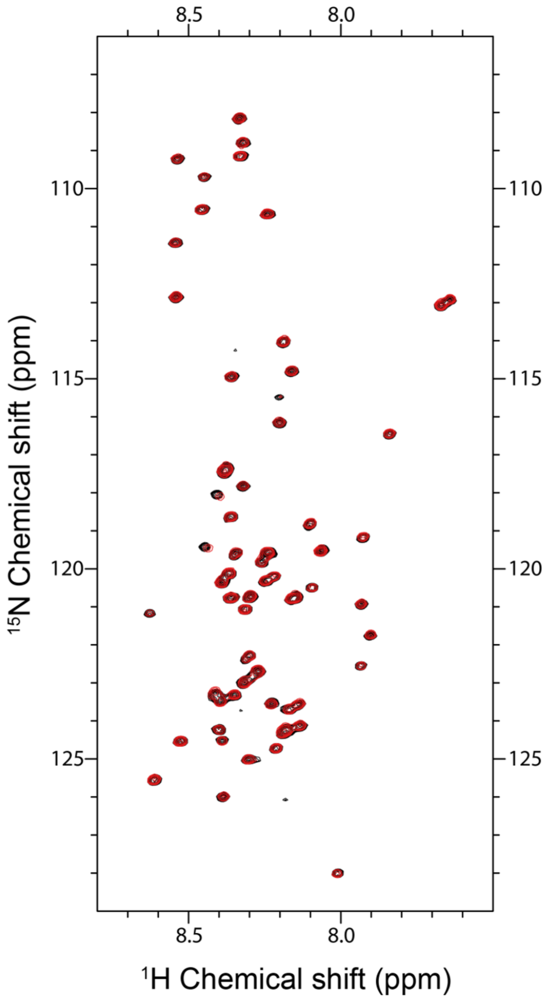


**Figure S13.** $^{1}$H-$^{15}$N HSQC spectra of 50 μM uniformly $^{13}$C-$^{15}$N labeled FtsZ C-tail in the absence (black) or presence (red) of 75 μM FtsQ1-99. Acquired at 800 MHz and 298 K in 20 mM phosphate buffer (pH 6.5) with 25 mM NaCl.

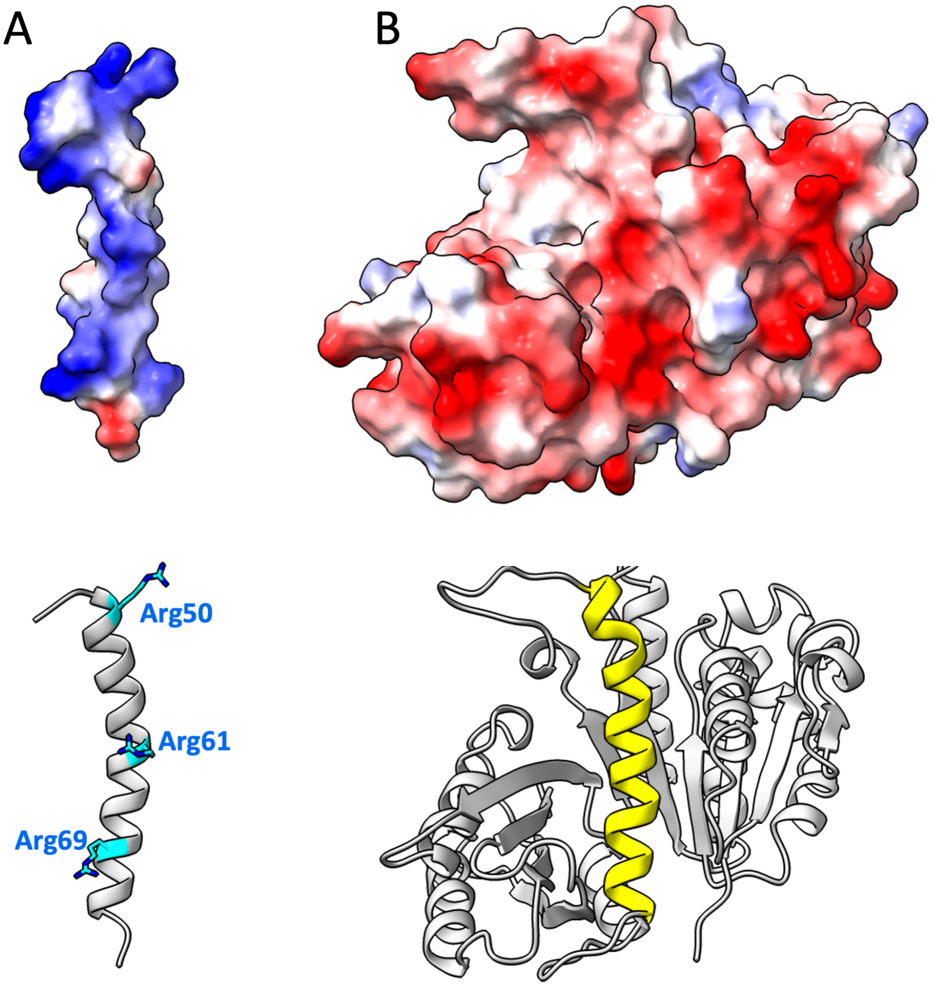


**Figure S14.** Electrostatic surfaces of (A) FtsQ Arg/Ala-rich helix and (B) FtsZ GTPase domain. As illustrated by the structures in the lower panels, the view of the FtsZ GTPase domain is directly into the inter-subdomain cleft (with the H7 helix at the base of the cleft in yellow); the view of the FtsQ helix is into the face that would be buried in the modeled complex shown in Fig. 8A. On this face are Arg 50, Arg61, and Arg69, shown as sticks.
